## Supplementary File 1 for "Structural foundation for the role of enterococcal PrgB in conjugation, biofilm formation and virulence"

| Strain, plasmid, or oligonucleotide | Relevant feature(s) or sequence | Source/reference or use |
| --- | --- | --- |
| **Strains** |  |  |
| *E. coli* |  |  |
| BL21 DE3 | Laboratory strain for expressing gene driven by T7 promoter | New England  Biolabs |
| Top10 | laboratory strain for cloning | Thermofisher Scientific |
| *E. faecalis* |  |  |
| OG1RF | donor strain, Rif^R^, Fus^R^ | (Dunny et al., 1981) |
| OG1ES | recipient strain, Ery^R^, Str^R^ | (Staddon et al., 2006) |
| **Plasmids** |  |  |
| pCF10 | cCF10-inducible conjugative plasmid, Tet^R^ | (Dunny et al., 1981) |
| pCF10Δ*prgB* | pCF10 plasmid with *prgB* deletion, originally designated as pCF10-8, Tet^R^ | (Chuang-Smith et al., 2010) |
| pET28-*prgB*_188-1233_ | N-terminal His tagged PrgB_188-1233_ expressing plasmid driven by T7 promoter, Kan^R^ | (Schmitt et al., 2018) |
| pET28-*prgB*_246-558_ | N-terminal His tagged PrgB _246-558_ expressing plasmid driven by T7 promoter, KanR | (Schmitt et al., 2018) |
| p7XC3GH-*prgB*_580-1233_ | C-terminal His and eGFP tagged PrgB_580-1233_ expressing plasmid driven by T7 promoter, KanR | This study |
| p7XC3GH-*prgB*_188-1233_ | C-terminal His and eGFP tagged PrgB_188-1233_ expressing plasmid driven by T7 promoter, KanR | This study |
| p7XC3GH-*prgB*_188-1233 S442A N444A_ | Mutant plasmid derived from p7XC3GH-*prgB*_188-1233_ | This study |
| pMSP3545S-*prgK* | Nisin-inducible shuttle plasmid expressing PrgK, SpcR, EryR | (Laverde Gomez et al., 2014) |
| pMSP3545S-MCS | pMPS3545S with additional NotI, BamHI, SphI, and SmaI sites, Spc^R^, Ery^R^ | This study |
| pMSP3545S-*prgB* | *prgB* coding sequence inserted between NcoI and BamHI in pMSP3545S-MCS | This study |
| pMSP3545S-*prgB*_ΔCOI_ | pMSP3545S-*prgB* with COI domain deletion | This study |
| pMSP3545S-*prgB*_ΔCSA1-CSC2_ | pMSP3545S-*prgB* with CSA1 domain to CSC2 domain deletion | This study |
| pMSP3545S-*prgB* _ΔCSA1-CSC1_ | pMSP3545S-*prgB* with CSA1 domain to CSC1 domain deletion | This study |
| pMSP3545S-*prgB* _ΔCSA2-CSC2_ | pMSP3545S-*prgB* with CSA2 domain to CSC2 domain deletion | This study |
| pMSP3545S-*prgB* _ΔCSA1_ | pMSP3545S-*prgB* with CSA1 domain deletion | This study |
| pMSP3545S-*prgB* _ΔCSC2_ | pMSP3545S-*prgB* with CSC2 domain deletion | This study |
| pMSP3545S-*prgB*_S442A_ | pMSP3545S-*prgB* with S442A point mutation | This study |
| pMSP3545S-*prgB*_S442A N444A_ | pMSP3545S-*prgB* with S442A N444A point mutation | This study |
| pMSP3545S-*prgB*_E455S_ | pMSP3545S-*prgB* with E455S point mutation | This study |
| pMSP3545S-*prgB*_S442A E455S_ | pMSP3545S-*prgB* with S442A E455S point mutation | This study |
| pMSP3545S-*prgB*_S442A N444A E455S_ | pMSP3545S-*prgB* with S442A N444A E455S point mutation | This study |
| **Primers** |  |  |
| MCS_fwd | catgggcggccgcggatccgcatgccccggggt | This study |
| MCS_rev | ctagaccccggggcatgcggatccgcggccgcc | This study |
| NcoI-prgB-F | ccggccatgggaatgaatcaacagactgaagtaa | This study |
| BamHI-stop-prgB-R | ccggggatccttattttgtttcttttctacgtttaaag | This study |
| prgB_COI delet_part_inv_F | ttctaaagat cataagaacgaaaacagctatgtcaatg | This study |
| prgB_COI delet_part_inv_R | cgttcttatg atctttagaagaaacgttgcctaagtc | This study |
| prgB_CSA1CSC2de_par_inv_F | gccaatgtc gaaaaaccacaaacaccaccag | This study |
| prgB_CSA1CSC2de_par_inv_R | ggtttttc gacattggctttgtatccattaaattc | This study |
| prgB_CSA1CSC1del_par_inv_F | gccaatgtc gatgatccaaaaccaaccaaagc | This study |
| prgB_CSA1CSC1del_par_inv_R | tggatcatc gacattggctttgtatccattaaattc | This study |
| prgB_CSA2CSC2del_par_inv_F | cacctgat gaaaaaccacaaacaccaccag | This study |
| prgB_CSAC2SC2del_par_inv_R | ggtttttc atcaggtgtatgcgtcaccac | This study |
| prgB_CSA1delet_part_inv_F | gccaatgtc agtagcaacccttccaaagacg | This study |
| prgB_CSA1delet_part_inv_R | ttgctact gacattggctttgtatccattaaattc | This study |
| prgB_CSC2 del_part_inv_F | attccaaaa gaaaaaccacaaacaccaccag | This study |
| prgB_CSC2 del_part_inv_R | ggtttttc ttttggaatatggttgacaacagtattg | This study |
| prgB_S442A_part_inv_F | tgtctgctttaaa ttcaagtttaacgaataaaggtg | This study |
| prgB_S442A_part_inv_R | tttaaagcagaca gcgcatacgcaaatggac | This study |
| prgB_S442AN444A_par_inv_F | tctgctttagcttca agtttaacgaataaaggtggc | This study |
| prgB_S442AN444A_par_inv_R | tgaagctaaagcaga cagcgcatacgcaaatgg | This study |
| prgB_E455S_part_inv_F | atgcgtcatttgt ttctgattttggggccaac | This study |
| prgB_E455S_part_inv_R | acaaatgacgcat ggccacctttattcgttaaac | This study |
| **DNA used for cryo-EM or EMSA** |  |  |
| 120 bp ssDNA | gcgaaatattggtaccccatggaatcgagggatcctctagtcgcaacatgct agcatgttgctccgcttgcaaaaagaaaagtcgacacgcgtagatctgctag catcgatccatggact | (Rehman et al., 2019) |
| DNA_100_ | tcgcaacatgctagcatgttgctccgcttgcaaaaagaaagcctacccttgggtata  accaattgtcaaactaaggagactacttattatgtaaaagaaa | (Schmitt et al., 2018) |
| PrgB_188-1233_ FX-F | atatatGCTCTTCtAGTgaAccgtaTgagaaagaagtcgcggaa | This study |
| PrgB_188-1233_ FX-R | tatataGCTCTTCaTGCaggcttcactgattgcgcatttaagtt | This study |
| PrgB_580-1233_ FX-F | atatatGCTCTTCtAGTgccaatgtcgttcctgttcttgttccg | This study |
