## Supplementary File 2 for "Structural foundation for the role of enterococcal PrgB in conjugation, biofilm formation and virulence"

| *Hardware* |  | | *Software* | |  |
| --- | --- | --- | --- | --- | --- |
| Microscope | Titan Krios | | **Data collection** | | EPU v 2.8.0 |
| Detector (mode) | K2 (counted) | | **Collection method** | | AFIS |
| Accelerating voltage | 300 keV | |  | |  |
| Spherical aberration | 2.7 | |  | |  |
| *Data acquisition parameters* |  | |  | |  |
| Apertures (C1, C2, C3) | 2000, 70, 2000 | | **Defocus range (µm, step size)** | | -1.2 to -2.7 (0.3) |
| Objective aperture | 100 | | **Dose (e/px/sec)** | | 7.75 |
| Energy filter slit (eV) | 20 | | **Dose (e/Å^2^/sec)** | | 11.5 |
| Illuminated area (µm) | 1.00 | | **Exposure time (sec)** | | 5 |
| Spot size | 6 | | **Total dose (e/Å^2^)** | | 57.5 |
| Tilt angle (°) | 0 | | **Dose fractions (#)** | | 40 |
| Nominal magnification | 165 000 x | | **Dose per fraction (e/Å^2^)** | | 1.44 |
| Pixel size (Å) | 0.82 | |  | |  |
| *Data processing parameters* | |  | |  | |
|  | | **PrgB apo** | | **PrgB ssDNA** | |
| Number of micrographs (total) | | 2797 | | 1670 | |
| Initial particle number (subset 500 micrographs) | | 170,826 | |  | |
| Initial particle number (total) | | 1,186,745 | | 714,102 | |
| Final particle number (total) | | 283,630 | | 163,566 | |
| Average map resolution (Å) | | 8-10 | | 9-11 | |
| FSC threshold | | 0.143 | | 0.143 | |
