## Supplementary File 3 for "Structural foundation for the role of enterococcal PrgB in conjugation, biofilm formation and virulence"

Table S3. Top hits of Dali search based on PDB of PrgB Ig-like domains

| No. | PDB id | Z score | rmsd | lali | nres | id (%) | description |
| --- | --- | --- | --- | --- | --- | --- | --- |
| 1 | 4ofq | 29.4 | 3.2 | 310 | 337 | 29 | AspA, 3 Ig-like domains, *S. pyogenes* |
| 2 | 4tsh | 28.1 | 4.8 | 432 | 485 | 25 | SpaP, 3 Ig-like domains, *S. mutans* |
| 3 | 4z1p | 26.8 | 3.3 | 298 | 334 | 21 | BspA, 2-Ig-like domains, *S. agalactiae* |
| 4 | 6e3f | 25.3 | 2.8 | 292 | 497 | 24 | Pas, 3 Ig-like domains, *S. intermedius* |
| 5 | 7l0o | 25.2 | 2.3 | 280 | 479 | 23 | SspB, 3 Ig-like domains, *S. gordonii* |
| 6 | 4igb | 15.0 | 3.5 | 159 | 435 | 13 | Sgo0707, 2-Ig-like domains, *S. gordonii* |
| 7 | 3kpt | 14.7 | 5.6 | 215 | 355 | 13 | BcpA, 3 Ig-like domains, *B. cereus* |
| 8 | 5xcc | 14.2 | 6.4 | 225 | 454 | 10 | CppA, 3 Ig-like domains, *C. perfringens* |
| 9 | 4hss | 13.6 | 5.1 | 209 | 411 | 16 | SpaD, 3 Ig-like domains, *C. diphtheriae* |
